# Evolution of SARS-CoV-2 reshapes upper airway immunity in children

**DOI:** 10.64898/2026.10.15.759133

**Authors:** Sophie McEwen, Jobst Röhmel, Sebastian Wiśniewski, Jennifer Loske, Marey Messingschlager, Johannes Liebig, Robert L. Chua, Anke Seegebarth, Foo Wei Ten, Birgit Sawitzki, Leif-Erik Sander, Christian Conrad, Roland Eils, Marcus A. Mall, Irina Lehmann, Saskia Trump

**Author notes:** Corresponding author: Prof. Dr. Irina Lehmann, Center of Digital Health, Molecular Epidemiology Unit, Berlin Institute of Health at Charité - Universitätsmedizin Berlin, Luisenstrasse 65, 10117 Berlin, mail. equal contribution (Marcus A. Mall, Irina Lehmann, Saskia Trump).

## Abstract

Children were relatively protected from severe COVID-19 during the early pandemic a phenomenon linked to rapid and heightened innate immune responses in the upper-airways¹,². Although adult upper-airway antiviral responses varied across successive SARS-CoV-2 waves³, whether viral evolution similarly reshaped pediatric mucosal immunity remains unclear. Here we used single-cell transcriptomics to profile 229,533 upper-airway cells from 68 unvaccinated children including 17 uninfected controls, spanning the wild-type SARS-CoV-2, Alpha, Delta and Omicron waves. Epithelial interferon responses were primarily determined by time since symptom onset but were, unlike in adults, independent of disease severity. During early infection, anti-viral responses differed across variants without following a progressive pattern of attenuation. Alpha and Delta elicited the strongest interferon programs across epithelial and immune cell populations, whereas Omicron exhibited a shift towards macrophage cytokine signaling and T-cell cytotoxicity. These findings reveal variant-specific remodeling of pediatric upper-airway immunity that fundamentally differs from that described in adults.

## Main Text

Previous studies identified enhanced epithelial viral sensing and a heightened innate and adaptive immune response in the childhood upper airway as characteristic features of the pediatric response to wild-type SARS-CoV-2, suggesting that these features may contribute to protection against severe COVID-19 ^1,4,5^. However, whether these distinctive features of pediatric immunity were maintained as SARS-CoV-2 evolved remains unknown. In adults, upper-airway interferon responses have been shown to vary with infecting variant and disease severity ^3^.

Here, we used single-cell RNA sequencing to profile nasal samples from 68 unvaccinated children, recruited in Germany between August 2020 and March 2022, including healthy controls, as confirmed by RT-PCR and clinical assessment (n=17), and children infected with wild-type SARS-CoV-2 (n=15) or the Alpha (n=7), Delta (n=12) or Omicron (n=17) strains. This enabled comparison of upper-airway immunity in unvaccinated children spanning four SARS-CoV-2 variant waves (*Fig. 1a, b, Supplementary Data Table 1*). Following quality control, we obtained 229,533 high-quality cells spanning the major epithelial and immune cell populations of the pediatric upper airways (*Fig. 1c; Extended Data Fig. 1)*.

**Figure 1:**
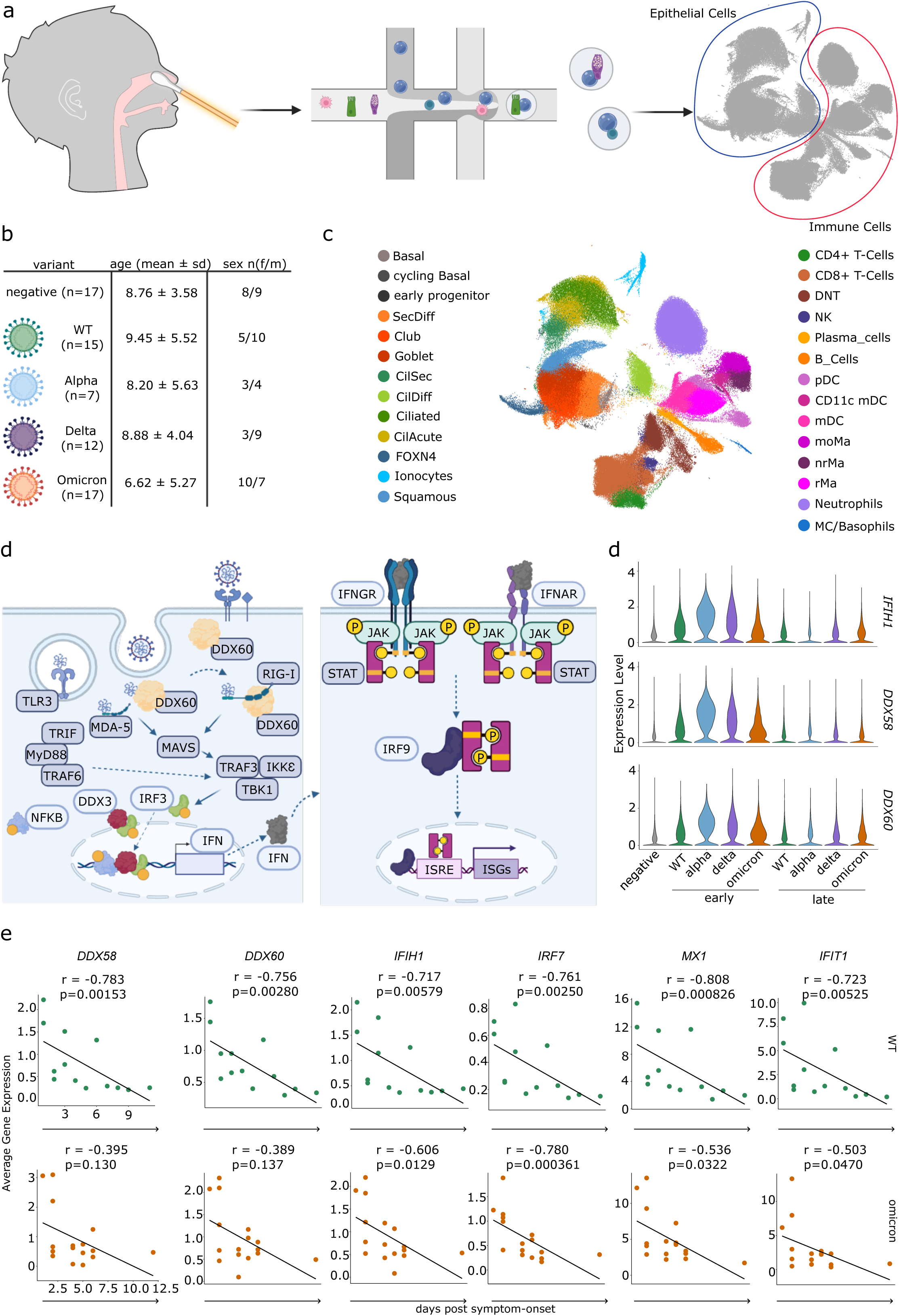
Antiviral Response is Strong during Early Infection. (a) Nasal swab samples were obtained from 68 pediatric participants for single cell RNA sequencing. (b) 17 participants were recruited as healthy controls. Fifty-one COVID-19 patients with either wild-type (n=15), Alpha (n=7), Delta (n=12) or Omicron (n=17) infections exhibited disease courses ranging from asymptomatic to severe were recruited. (c) Uniform Manifold Approximation and Projection (UMAP) of all 68 samples showing immune and epithelial cell types identified in nasal swab data set. (d) Initial stages of viral detection and antiviral response activation upon SARS-CoV-2 infection. (e) Mean of normalized expression across epithelial cells, shown for pattern recognition receptors per condition in early (≤4 days post symptom-onset) or late (>4 days post symptom-onset) stages of infection. Analysis excluded asymptomatic patients or those without dps information (n=64). (f) Per sample mean of normalized expression across epithelial cells for antiviral genes, shown in relation to the number of days since symptom-onset at sampling in wild-type (n=12) and Omicron (n=15). Asymptomatic patients or those without dps information excluded (n=64).

We first examined SARS-CoV-2 sensing and downstream antiviral responses across the epithelial compartment, focusing on key pattern recognition receptors (PRRs) and downstream signaling mediators across infection phases, as outlined schematically in *Fig. 1d*. Based on our previous study, which identified a change in the expression dynamics of the SARS-CoV-2 pattern recognition receptors (PRRs) *IFIH1* (MDA-5) and *DDX58* (RIG-I) around day 4 post symptom onset ^1^, we stratified our samples into early (≤4 days post symptom (dps) onset) and late (>4 dps) infection. Across variants, epithelial expression of *IFIH1, DDX58,* and *DDX60* was highest during early infection and declined during the late phase towards levels observed in healthy controls (*Fig. 1e*).

To further resolve the temporal dynamics of epithelial antiviral responses, we correlated the expression of *DDX58, DDX60, IFIH1, IRF7, MX1* and *IFIT1* with dps-onset in children infected with wild-type SARS-CoV-2 or Omicron, for which samples covered a broad dps range. In wild-type SARS-CoV-2 infection, all six genes showed strong negative correlations with time since symptom onset (Spearman’s r = -0.81 to -0.72, all p-value < 0.01). This is consistent with human SARS-CoV-2 challenge studies, showing an early rise followed by a decline of nasopharyngeal interferon responses after infection with a wild-type strain^6^. Correlations were generally weaker during Omicron infection (Spearman’s r = -0.78 to -0.50, all p-value < 0.05) and were no longer significant for *DDX58* and *DDX60* (*Fig. 1f*). These findings support a focus on early infection when comparing variant-associated epithelial antiviral responses and suggest differences in their temporal regulation across variants.

To characterize the upper-airway epithelial response beyond selected antiviral mediators, we performed pathway enrichment analysis on the top 200 differentially expressed genes (DEGs) between healthy controls and all SARS-CoV-2+ samples (*Supplementary Data Table 2*). Enriched pathways were associated with antiviral response programs and to a lesser extent, cilium movement (*Extended Data Fig. 2a)*. Analysis of genes contributing to these pathways across variants and infection phases revealed three distinct clusters (*Fig. 2a*). The largest, *Cluster 1,* comprised PRRs, downstream signaling mediators, and interferon-stimulated genes (ISGs). Conditions separated primarily by infection phase, with late-phase responses clustering with healthy controls. Within early infection, however, marked variant-associated differences emerged: Alpha and Delta induced the strongest expression of antiviral response genes, while wild-type and Omicron elicited a more moderate response. Notably, the strong early antiviral response to Alpha coincided with induction of cilium-associated gene expression, linking variant-specific antiviral activation to broader changes in the transcriptional program of the upper airway epithelium.

**Figure 2:**
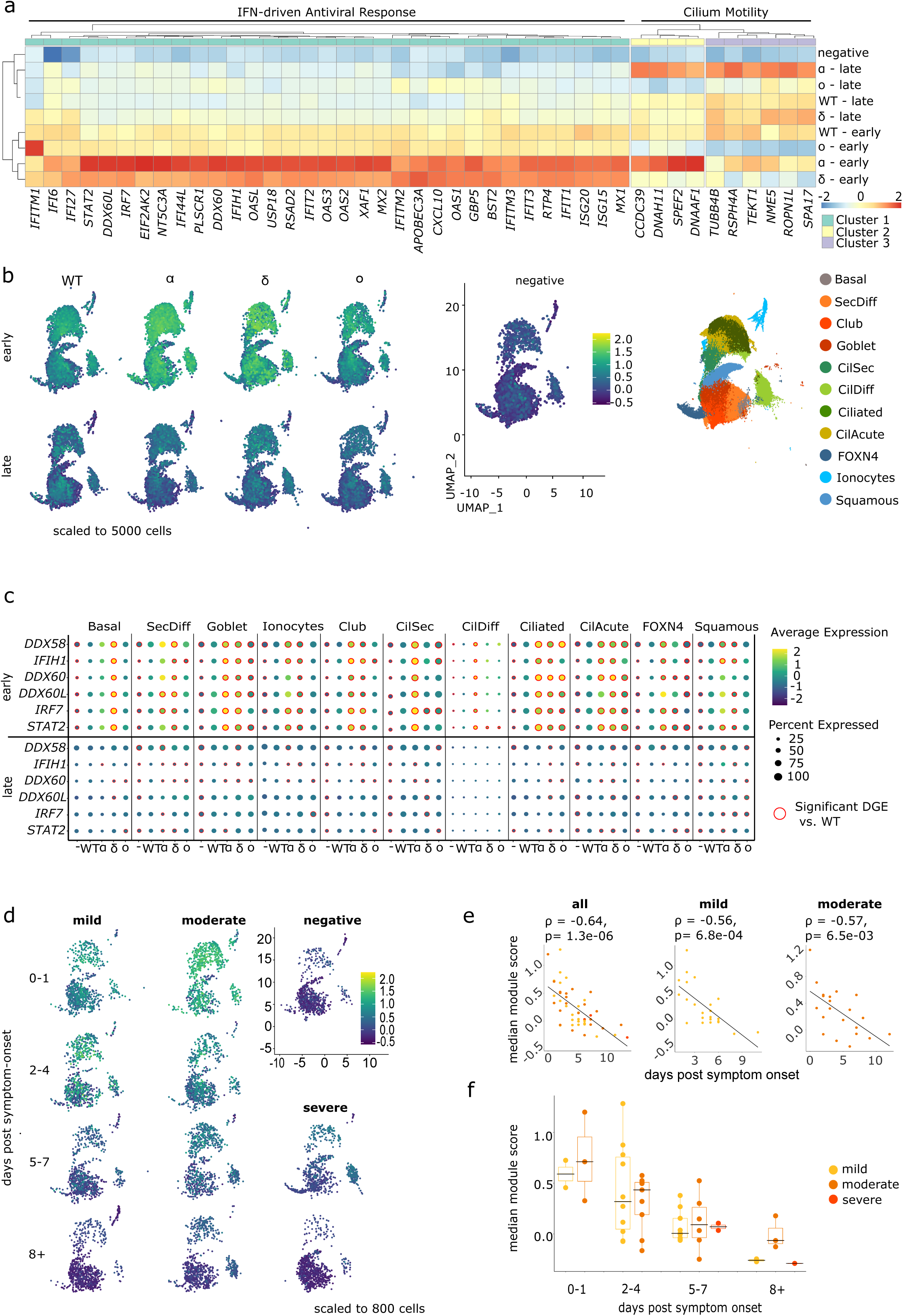
Infecting Variant and Infection Stage Trump Disease Severity when Determining Magnitude of Epithelial Antiviral Response. Asymptomatic samples and positive samples without dps information were excluded (n=64). (a) Mean normalized expression of genes from the top 10 enriched Gene Ontology terms among the top 200 differentially expressed genes between healthy controls and SARS-CoV-2+ samples (fdr < 0.05 & |log2FC| > 1), shown across conditions and infection-stages. (b) Per cell module scores for antiviral response genes (Cluster 1) shown across infection-stage and condition in epithelium. Each plot down sampled to 5000 cells. (c) Mean normalized expression for antiviral response genes, shown across epithelial cell types, per condition and infection stage. Significant differential expression vs. wild-type is indicated by a red circle (fdr < 0.05 & |log2FC| > 1). (d) Per cell module scores of genes from Cluster 1 of Figure 2a in the epithelium across severity categories and number of days post symptom-onset (dps). All plots were down sampled to 800 cells. (e) Median module score per sample vs. dps. Spearman’s Rho (ρ) and P-value reported (all [mild, moderate, and severe]: ρ = -0.64, p = 1.3 x 10-06; mild: ρ = -0.56, p = 6.8 x 10-04; moderate: ρ = -0.57, p = 6.5 x 10-03). (f) Median module score per sample across dps and severity categories.

Genes involved in axonemal dynein assembly and motility (*CCDC39, DNAH11, SPEF2, DNAAF1*) were particularly strongly induced following Alpha infection, while a second cluster containing genes encoding structural and radial-spoke components (*TUBB4B, TEKT1, RSPH4A, NME5*) was induced during late Alpha and Delta infection^7,8^. This induction of ciliary transcriptional programs in children, especially following Alpha infection, stands in contrast to previous studies in adults. Adults infected with wild-type SARS-CoV-2 exhibited repression of cilium-associated genes during acute infection which persisted after viral clearance^9^. As ciliated cells are a major target of SARS-CoV-2, the concurrent induction of cilia-associated and antiviral programs may reflect better preservation of epithelial integrity in children, potentially supported by the pre-activated antiviral state previously described in the pediatric upper-airway 1,5. Consistent with this interpretation, Alpha infection has been associated with less pronounced ciliary loss in pediatric than in adult upper-airway epithelial cultures ^10^.

To determine whether the *Cluster 1* antiviral response was restricted to specific epithelial cell types or states, we calculated a module score for each epithelial cell based on all genes in this cluster. Antiviral activation was broadly distributed across multiple epithelial populations, without clear restriction to a specific cell type or state (Fig. 2b). Variant-associated differences were most pronounced during early Alpha and Delta infection and largely diminished during the late phase (*Fig. 2b*). Consistently, key PRRs (*DDX58, IFIH1, DDX60*) and downstream mediators (*DDX60L, IRF7, STAT2*) exhibited similar condition-dependent expression patterns across multiple epithelial populations (*Fig. 2c, Extended Data Fig. 2b, Supplementary Data Table 3*). Several epithelial cell-types showed significantly higher expression in early Alpha and Delta than wild-type infection (*Fig. 2c, Supplementary Data Table 3*). These findings indicate that the heightened Alpha/Delta interferon response is an epithelium-wide state rather than the consequence of a single highly responsive cell population. In contrast to findings in adults, the epithelial responses in children were similarly pronounced in Alpha and Delta, while Omicron more closely resembled the more moderate response elicited by the wild-type.

In adults, upper-airway ciliated cell interferon responses following wild-type and Delta infection have been inversely associated with disease severity, an association not observed in Omicron ^3^. In our pediatric cohort, however, the magnitude of the interferon response was primarily structured by time since symptom onset rather than disease severity. *Cluster 1* module scores declined similarly over time across severity categories (*Fig. 2d*). Sample-level scores correlated negatively with days post symptom onset across the cohort (Spearman’s r*= -0.64, p=1.3*10^-6^, Fig. 2e*) and within mild (Spearman’s r*= -0.56, p=6.8*10^-4^*) and moderate (Spearman’s r*= -0.57, p=6.5*10^-3^*) disease. Accordingly, no significant differences in module scores were observed between severity groups within days post symptom bins (*Fig. 2f*). Thus, within the predominantly mild-to-moderate pediatric disease spectrum studied here, infection stage rather than severity was the major determinant of epithelial interferon activity.

In vitro studies have demonstrated enhanced antagonism of innate immune responses by certain SARS-CoV-2 variants, including Alpha and the Omicron sub-lineages BA.4 and BA.5. However, in our primary pediatric samples, Alpha induced one of the most pronounced innate antiviral responses, highlighting that virus-intrinsic immune evasion does not necessarily translate into attenuated antiviral responses within the complex multicellular environment of the upper-airway epithelium. In contrast, the limited epithelial interferon response to Omicron may reflect variant-specific differences in age-dependent host–virus interactions. Consistent with this possibility, previous studies have shown that the reduced viral replication observed in pediatric versus adult upper-airway epithelial cells following infection with ancestral SARS-CoV-2 or Delta was less pronounced in Omicron¹³.

An important limitation of our study is that variant waves were inseparable from calendar time, and we therefore cannot exclude undocumented prior SARS-CoV-2 infections as a contributor to the observed variant-associated differences, particularly for later infections. However, previous studies comparing vaccinated and unvaccinated individuals suggest that pre-existing immunity may enhance early upper-airway interferon responses upon subsequent SARS-CoV- 2 infection, rather than suppressing them ^14^.

Effective viral control requires coordination between the epithelial antiviral response and immune-cell recruitment. Using CellChat, we investigated intercellular communication to determine whether differences between variants extend beyond epithelial-intrinsic responses to changes in communication across the immune and epithelial compartments (*Supplementary Data Table 4*) ^15^. *MHC-I*, *MHC-II* and *Midkine* (*MK*) were among the most strongly interacting pathways across all conditions but varied little between variants (*Extended Data Fig. 3a*). In contrast, *CCL* and *ANNEXIN* signaling showed the most pronounced variant-dependent differences among the top 10 pathways identified for each variant (*Extended Data Fig. 3b*).

Macrophages were a major recipient population in both pathways, but their dominant cellular sources differed. *CCL* signaling was largely immune-derived, and strongest in Alpha and Delta, whereas *ANNEXIN* signaling was primarily epithelium-derived and targeted macrophages, particularly in Delta and Omicron (*Fig. 3a*). This communication was dominated by *ANXA1– FPR1* interactions (*Supplementary Data Table 4; Extended Data Fig. 3c*) and accompanied by increased macrophage *FPR1* and, to a lesser extent, *FPR2* expression (*Extended Data Fig. 3d; Supplementary Data Table 5*). While *FPR1* promotes macrophage chemotaxis and antimicrobial effector functions, *ANXA1–FPR2* can exert immunoregulatory and pro-resolving effects^16,17^. Neither receptor differed between healthy controls and wild-type infection, but both were significantly increased in Alpha, Delta, and Omicron (*Extended Data Fig. 3d; Supplementary Data Table 5*). *ANXA1–FPR2* interactions emerged from Alpha onwards, potentially providing a counter-regulatory signal alongside inflammatory *CCL* communication^18^.

**Figure 3:**
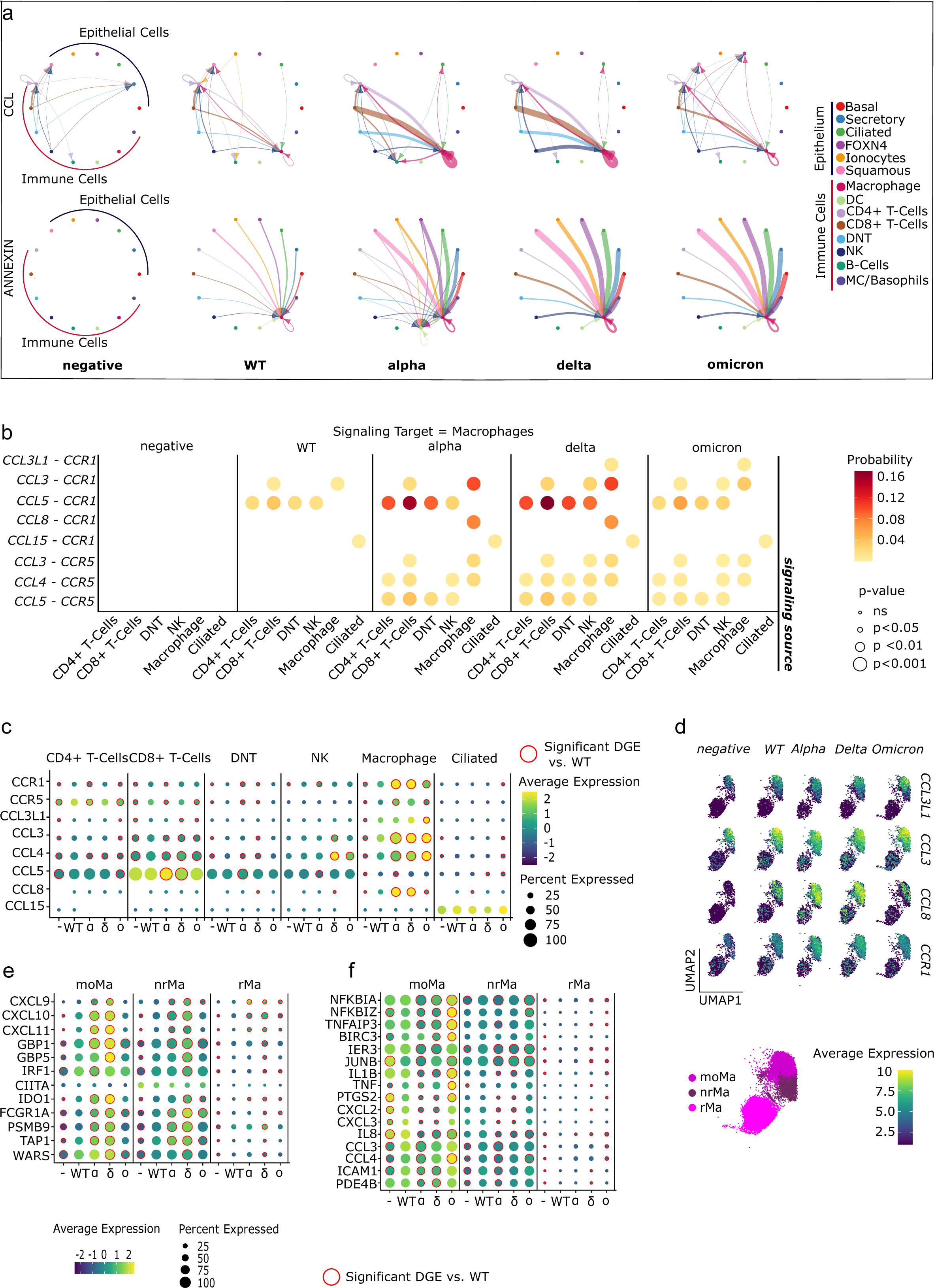
Variant-Dependent Differences in Macrophage Interactions and Activity. Analyses include controls, as well as all mild and moderate COVID-19 samples (n=62). (a) Interactions between bulk cell types for CellChat pathways *CCL* and *ANNEXIN* per variant. Probability is represented by line thickness, and arrowheads point towards target cell type. All shown interactions are significant (p<0.05). Interaction probabilities are scaled by maximum interaction weight per pathway. (b) Interaction probabilities for significant ligand-receptor pairs involved in *CCL* pathway interactions (p<0.01) where macrophages are the target (receptor-expressing) cell type. Source (ligand-expressing) cell types are shown on the x-axis. (c) Mean normalized gene expression levels for *CCL* ligands and receptors in interacting cell types. Average expression is shown per condition. A red circle indicates significance vs. wild-type (fdr < 0.05 & |log2FC| > 1). (d) Feature plot of macrophage subtypes showing average expression per cell of *CCL3L1*, *CCL3*, *CCL8* and *CCR1* across conditions. Each feature plot is scaled to 2300 cells. Mean normalized gene expression per condition of (e) interferon response genes and (f) cytokine response genes across macrophage subpopulations. A red circle indicates significance vs. wild-type (fdr < 0.05 & |log2FC| > 1).

We next resolved the macrophage-directed CCL interactions (*Fig. 3b*). Macrophage-derived *CCL3*-*CCR1* and *CCL8-CCR1,* as well as T-cell derived *CCL5-CCR1* interactions were particularly prominent in Alpha and Delta. Ciliated epithelial cells also contributed inferred *CCL15–CCR1* interactions, suggesting convergence of macrophage-, lymphocyte- and epithelial-derived CCL signals on macrophages (*Fig. 3b*). Consistent with these patterns, macrophage *CCR1* and *CCL8* expression were higher in Alpha and Delta than in wild-type infection, whereas *CCL3* expression increased in Alpha, Delta, and Omicron (*Fig. 3c, Supplementary Data Table 6*). Increased *CCL5–CCR1* interaction probabilities involved both CD4+ and CD8+ T cells in Alpha and Delta, although significant induction of *CCL5* itself was confined to CD8+ T cells (*Fig. 3c; Supplementary Data Table 6*). These findings point to CCR1- expressing macrophages as a convergence point for variant-dependent chemokine signaling.

The CCL ligands and receptors underlying these interactions were enriched in recruited non-resident (nrMa) and monocytic (moMa) macrophages, which showed increased *CCR1* and *CCL8* expression, in Alpha and Delta (*Fig. 3d*). Macrophage transcriptional states mirrored the variant-dependent epithelial response. Alpha and Delta were associated with stronger interferon-response signatures, including *GBP1*, *GBP5*, *CXCL10* and *CXCL11¹⁹˒²⁰* (*Fig. 3e; Supplementary Data Table 7*), while in Omicron, macrophages expressed NF-kB-associated cytokine genes such as *TNF, IL1B* and *CCL4*^21–23^ (*Fig. 3f; Supplementary Data Table 7*). Thus, Alpha and Delta elicited coordinated epithelial and macrophage interferon responses, whereas Omicron was associated with a more cytokine-dominant macrophage state. These differences were most pronounced during early infection and coincided with variant-dependent epithelial-to-macrophage ANNEXIN signaling, potentially providing counter-regulatory input to macrophage activation (*Extended Data Fig. 3e, 3f; Supplementary Data Table 5*).

*CCR1*-mediated macrophage recruitment to the upper airway has been associated with the pronounced inflammatory response observed in adults with severe or critical COVID-19^24,25^. Our findings suggest that recruited macrophages are also central to variant-specific differences in the immune cell responses to SARS-CoV-2 in children. As IFN-γ can induce CCR1 expression in monocytes^26^, this pathway may link the interferon-rich environment elicited by Alpha and Delta to enhanced CCR1-dependent communication. However, because our cohort predominantly comprised children with mild and moderate disease, these data do not establish whether increased *CCR1*-associated signaling contributes to immunopathology or supports an effective pediatric antiviral response.

Having examined lymphocytes as a source of CCL signaling, we next investigated variant-associated changes in T-cell-intrinsic transcriptional programs. We identified the top 200 cell-type specific DEGs between healthy controls and all SARS-CoV-2+ samples for CD8+ T-cells, CD4+ T-cells and NK cells (*Supplementary Data Table 8; Supplementary Data Table 9)*. Network-driven functional enrichment analysis of CD8+ and CD4+ T-cell DEGs identified seven Gene Ontology Biological Process modules, including two *Interferon Response* modules, a *Cell Death Mediators* module and an *Inflammatory Response/Response to Hypoxia* module (*Fig. 4a*, *Extended Data Fig. 4a*, *Supplementary Data Table 10*)^27^. Variant-associated differences were concentrated in the two *Interferon Response* and the *Cell Death Mediators* modules *(Fig. 4b; Extended Data Fig. 4b).* Given the broadly concordant patterns between CD4+ and CD8+ T-cells, subsequent gene-level analyses focused on the more abundant CD8+ T-cell populations; corresponding DEGs are listed in *Supplementary Data Table 8*.

**Figure 4:**
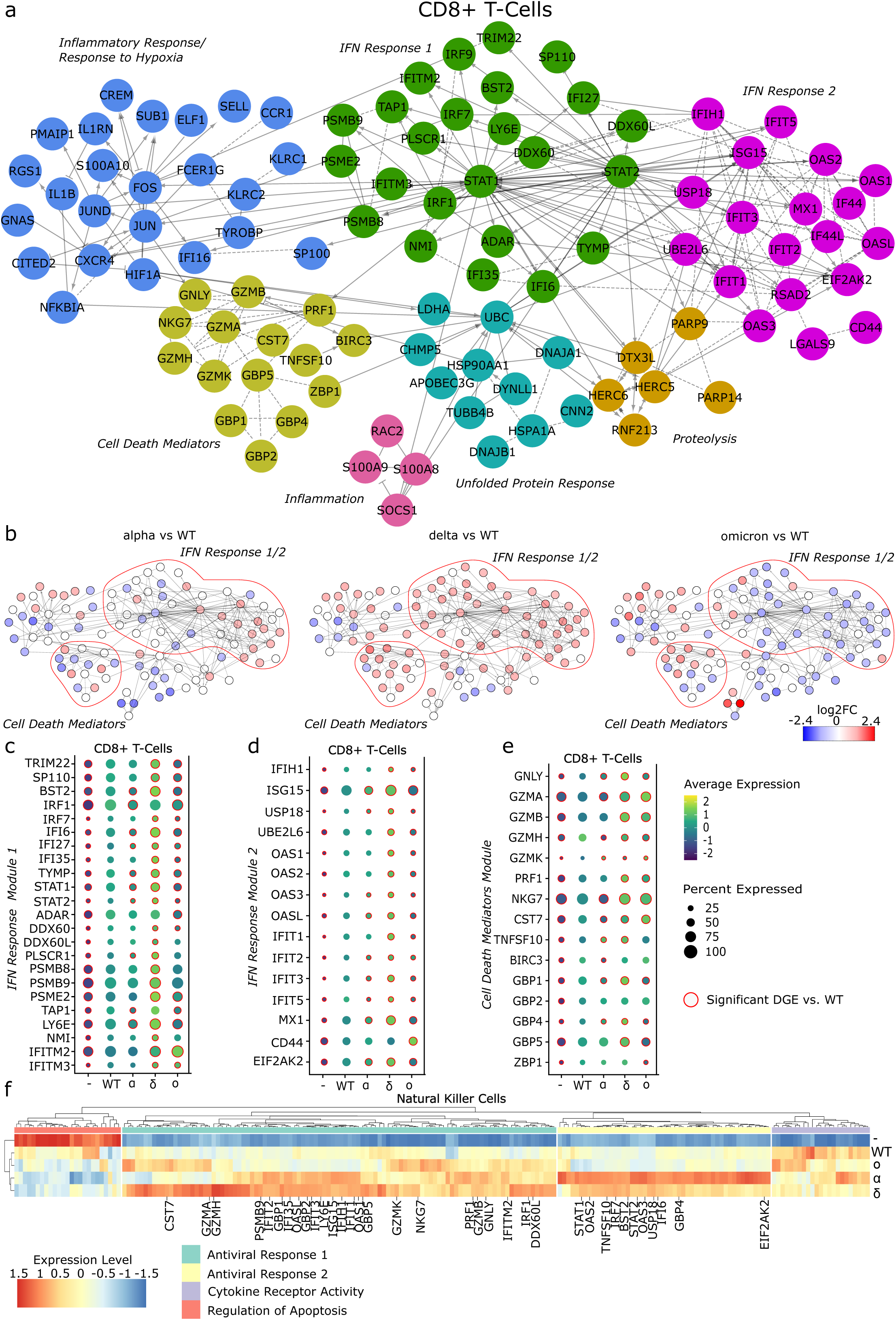
Divergent IFN Response and Cell Death Mediation between Variants in NK and CD8+ T-Cells. Analyses include controls, as well as all mild and moderate COVID-19 samples (n=62). Cytoscape networks based on top 200 differentially regulated genes (fdr < 0.05 & |log2FC| > 1) between healthy controls and SARS-CoV-2+ samples in CD8+ T-cells. Node color represents (a) module membership (b) log_2_FC for each variant vs. wild-type (fdr > 0.05). Comparison of average expression between variants (c) *IFN Response 1* module (d) *IFN Response 2* module and the (e) *Cell Death Mediators* module in CD8+ T-Cells. A red circle indicates significance vs. wild-type (fdr < 0.05 & |log2FC| > 1). (f) Mean normalized gene expression for top 200 differentially expressed genes (fdr < 0.05) between healthy controls and all SARS-CoV-2 positive samples in NK cells. Genes shown if maximum expression level/minimum expression level > 1.

At the gene level, both *Interferon Response* modules were coordinately upregulated in Alpha and Delta (*Fig. 4c, d*), extending the interferon-rich state observed in epithelial cells and macrophages to the lymphocyte compartment. *Cell Death Mediators* were most strongly affected in Delta and Omicron (*Fig. 4e*) but differed in composition: Omicron was dominated by cytotoxic effectors such as *GZMA, GZMB* and *PRF1*, whereas Delta additionally featured interferon-induced mediators including *GBP1, GBP4* and *GBP5* (*Fig. 4b, c*). Interferon modules were expressed most strongly during early Alpha and Delta infection and declined later (*Extended Data Fig. 4c*). In contrast, Omicron was associated with a sustained canonical cytotoxic CD8+ T-cell program, despite an interferon response largely comparable to healthy controls.

In NK cells, clustering of the top 200 infection-associated DEGs identified four clusters: two *Antiviral Response* clusters, a *Cytokine Receptor Activity* cluster, and a *Regulation of Apoptosis* cluster (*Fig. 4f; Supplementary Data Table 10*). As in epithelial cells, macrophages and T cells, antiviral gene expression was strongest in Alpha and Delta, including interferon-responsive and cytotoxic effector genes such as *GZMB, PRF1* and *GNLY*, whereas the Omicron-induced response more closely resembled the wild-type. Our lymphocyte analyses showed that the Alpha/Delta interferon-rich state extended across multiple epithelial and immune cell types. Notably, while interferon signaling was attenuated in Omicron, the cytotoxic effector response was upregulated compared to earlier variants, suggesting remodeling rather than uniform suppression of the antiviral immune response with viral evolution.

Together, our findings extend previous observations of enhanced pediatric upper-airway antiviral immunity to wild-type SARS-CoV-2 by demonstrating that these responses are shaped by both infection-stage and viral variant. In contrast to reports in adults, epithelial interferon activity in children is primarily associated with time since symptom onset rather than disease severity. Moreover, ciliary transcriptional programs were maintained or induced rather than suppressed, potentially reflecting better preservation of epithelial integrity, promotion of differentiation into ciliated cells, and improved muco-ciliary clearance. SARS-CoV-2 evolution remodeled pediatric upper airway immunity: Alpha and Delta elicited coordinated interferon-rich responses across epithelial and immune compartments, whereas Omicron shifted towards cytokine and cytotoxic programs. This variant-specific pattern is distinct from that reported in adults, where Delta and Omicron showed more similar response levels. Thus, our study provides evidence that temporal dynamics and variant dependence of antiviral mucosal immunity differ between children and adults, underscoring the importance of studying pediatric antiviral responses independently, rather than extrapolating from adult immunity.

## Methods

### Patient Recruitment and Ethical Approval

Pediatric study participants were recruited in Berlin, as part of the observational study, increased Resilience of Children compared to Adults in SARS-CoV-2 infection (RECAST), an arm of the prospective observational cohort study Pa-COVID-19. Recruitment took place between the 11^th^ of August 2020 and 31^st^ of March 2022, and covered healthy controls, as well as children infected with one of four variants of concern (ancestral/wild-type (WT), Alpha, Delta and Omicron), as defined by the World Health Organization (WHO), that circulated widely in Europe during that time period^28^. Recruited patients were categorized as either asymptomatic, mild, moderate, or severe, according to WHO guidelines^29^. Sixty-eight participants between the ages of 0 and 17 were recruited, see *Supplementary Data Table 1*. Sample size was determined by sample availability.

Healthy control donors were recruited between 7^th^ October 2020 and 9^th^ December 2020. Patients with a SARS-CoV-2 infection were recruited during the periods in which the respective variants were dominant in Germany: ancestral strain (WT), 11^th^ August 2020 – 5^th^ January 2021; Alpha, 26^th^ February 2021 – 28^th^ May 2021; Delta, 20^th^ August 2021 – 26^th^ November 2021; Omicron BA.1/BA.2, 20^th^ January 2022 – 31^st^ March 2022). The absence of SARS-CoV-2 infection was confirmed by a negative PCR test for SARS-CoV-2 Envelope (E) and Nucleocapsid (N), as described in Loske et al ^1^.

All participants were unvaccinated and presented without respiratory co-morbidities or previous SARS-CoV-2 infection, according to parental and/or participant report. Written, informed consent was obtained on behalf of all participants. All samples were collected in line with the Declaration of Helsinki and approval for the study was granted by the Institutional Review Boards (Pa-COVID-19/RECAST: EA2/066/20, SC2: 123/20-ek). Sample size was determined by participants’ availability.

### Isolation of Single Cells and Preparation of Libraries from Nasal Swabs

For all study participants, swabs of the nasal turbinate were obtained and processed according to previously described processes ^1, 9, 24, 25^,

Upon sampling, swabs were immersed in 500µL of precooled DMEM/F12 medium (Gibco). Samples were stored on ice, never exceeding the duration of one hour. Subsequently, 500µL of Dithiothreitol (AppliChem) was added to the swab to facilitate mucus disintegration. Gentle pipette-mixing of the suspension against the swab and 20 repetitions of submersion and lifting using tweezers, served to dislodge cells. Samples were incubated in a thermomixer (37 *°C*, 500rpm, 10 min). Centrifugation (300xg, 4*°C*, 5 min) and supernatant removal followed. In the case of red coloration of the pellet, treatment with Red Blood Cell Lysis Buffer (Roche, 11814389001) was applied (25 *°C,* 10 min), followed by centrifugation (300xg, 4*°C*, 5 min).

For logistical reasons, some samples were transferred to storage at -80 *°C* in cryopreservation medium [70% DMEM/F12 (Gibco), 20% fetal bovine serum (FBS, Gibco), 10% Dimethyl sulfoxide (Sigma-Aldrich)]. At the point of further use, thawing at 37 *°C* and centrifugation (300xg, 4*°C*, 5 min) preceded continuation of the protocol.

All samples were treated with 500µL Accutase (Thermo Fisher) for 10 minutes at room temperature. Enzymatic digestion was terminated by addition of 500µL of medium (DMEM/F12, 10% FBS), followed by centrifugation (300xg, 4*°C*, 5 min). Cells were resuspended in 100 µL PBS and filtered through a 35µm cell strainer (Falcon).

Libraries were prepared from 17,500 cells per sample using Chromium Next GEM Single Cell 3’ Kits v3.1 (10X Genomics) with all steps proceeding according to manufacturer’s protocol. An incubation step at 85 *°C* was added to the GEM-RT incubation step to ensure inactivation of viral particles.

Final libraries were sequenced using the NovaSeq 6000 Sequencing System (Illumina; paired-end, single index).

### Data Processing and Quality Control

FASTQ files were processed using CellRanger version 3.1.0 (10X Genomics). Transcripts were aligned against a custom GRCh37/hg19 genome assembly with the SARS-CoV-2 genome appended as a single gene (10X Genomics, RefSeq ID: NC_045512). Quality control steps were carried out in Seurat version 4.3.0, running on R version 4.0.5^30^. Cells with a mitochondrial read proportion of 15% or higher or expressing fewer than 200 genes were removed from each sample. Samples with a cellular detection rate lower than 1000 were removed from the dataset (*Supplementary Table 11*). For each sample, setting an individual upper cutoff for UMI count, based on a scatter plot of gene count vs. UMI count, served as a manual method for doublet removal.

Samples were imported into Scanpy 1.9.1, using AnnData 0.7.5 and running on python version 3.10.531. Reads were normalized to 10,000 per cell and expression values were log-transformed and scaled. The following pre-processing steps were conducted on the 5000 most highly variable genes. Data integration was performed on PCA transformed data, using Harmony version 0.0.5 (theta = 2, sigma = 0.1, lamb = 1, block_size = 0.0025) and BBKNN version 1.5.1 (batch_key = “sample”, trim = 100) ^32,33^. UMAP embedding and Leiden clustering followed. For finer definition of variable cell types, such as Macrophages, subsets were re-clustered at higher resolution, also using the Leiden algorithm. Immune and epithelial cell types were annotated manually using common marker genes (*Extended Data Fig. 1*). The annotated object was converted and analyzed further in R 4.0.5 using Seurat 4.3.0.

### Data Analysis and Statistical Testing

Differential gene expression (DGE) analyses were conducted in Seurat, using the MAST model, with age and sex of the donor, in addition to the fresh/frozen status of the sample, considered as covariates across all experiments. The Benjamini-Hochberg method for multiple testing correction was applied. Significance was determined if the false discovery rate (FDR) was smaller than 0.05 and the log_2_FC was greater than or equal to 0.1 or smaller than or equal to –0.1. Cell types consisting of fewer than 200 cells across all samples were excluded from DGE analyses. Mitochondrial-encoded (MT-) and ribosomal protein (RP-) genes were removed prior to calculation.

Module scores, which represent average expression levels of chosen gene sets per individual cell, with consideration to background expression, were calculated using the *AddModuleScore* function in Seurat. All conditions were downsampled to a maximum number of cells per identity when calculating a module score or representing the module score as a UMAP. This was done using the *downsample* parameter within the *subset* function. For the epithelial antiviral gene expression module, a Spearman’s Rank Correlation Coefficient was calculated to represent the relationship between the median module score per sample and days post symptom onset.

CellChat v1 ^15^ was used for investigation of cell-cell interactions. Cell type bulks were used for CellChat Analyses. Cell type bulk categories were defined as follows: Basal = [early progenitor, cycling_Basal, Basal]; Ciliated = [CilSec, CilDiff, CilAcute, Ciliated]; Secretory = [SecDiff, Club, Goblet]; Squamous = [Squamous]; FOXN4 = [FOXN4]; Ionocytes = [Ionocytes]; Macrophage = [moMa, nrMa, rMa]; MC/Basophils = [MC/Basophils]; CD4+ T-Cells = [CD4+ T-Cells]; CD8+ T-Cells = [CD8+ T-Cells]; NK = [NK]; DNT = [DNT]; B-Cells = [Plasma Cells, B Cells]; DC = [mDC, pDC, CD11c+ mDC]. Neutrophils were excluded on the grounds of their sensitivity to freezing/thawing cycles, and severe patients were excluded from immune and cell-cell interaction analyses.

Interactions were calculated per condition, based on the human CellChat database. No protein-protein interaction projection was used. Communication probability was calculated based on raw data. Interactions present in at least 10 cells and with a p-value of 0.05 or less were included in the final interaction network.

To select pathways of interest, a mean probability score was calculated per pathway. Ten pathways with the highest probability of interaction were selected for each condition. Pathways that exhibited interactions in all conditions were removed to draw focus to high-probability pathway interactions; that were also highly variable between conditions. A heatmap was plotted, pathways were clustered by mean probability score, using the Ward.D2 algorithm. The two pathways with the highest mean probability score across conditions were visually selected from the heatmap for further investigation (*Extended Data Fig. 2a*).

Circle plots were generated per condition, per pathway, with edges representing total communication probability between cell types. Plots were scaled by maximum edge weight across conditions, per pathway. All interactions where p < 0.05 were shown. All ligand-receptor interactions of a pathway of interest were considered for further analysis.

Cytoscape networks were generated using Cytoscape version 3.8.2 in combination with the *GeneSet/Mutation Analysis* tool from the ReactomeFIPlugIn version 2020 ^27^. Networks were constructed based on the top 200 most strongly differentially regulated genes between healthy controls and all SARS-CoV-2 positive samples, determined by absolute log_2_FC. No linker genes were used. Following clustering, modules were annotated based on significantly enriched terms of the Gene Ontology Biological Process. These were compared between conditions, based on the log_2_FC between genes in WT and each variant individually. Modules containing fewer than three genes were removed.

Gene Set Enrichment Analysis, using *EnrichR* v3.4, was carried out to annotate heatmap clusters. Input gene lists, in this case genes present in each cluster, were tested against the *Gene Ontology Molecular Function 2023* and *Gene Ontology Biological Process 2023* databases and clusters annotated based on top 10 enriched terms by p-value. All input genes were significantly differentially expressed (fdr 0.05) with a log_2_FC < -0.1| > 0.1.

## Acknowledgements

We thank all patients of the RECAST and Pa-COVID-19 studies for donating nasal samples and clinical data. This study was supported by the BIH COVID-19 research program and the fightCOVID@DKFZ initiative, the European Commission (ESPACE, 874719, Horizon 2020), the German Federal Ministry of Education and Research (BMBF)-funded de.NBI Cloud within the German Network for Bioinformatics Infrastructure (de.NBI; 031A537B, 031A533A, 031A538A, 031A533B, 031A535A, 031A537C, 031A534A, and 031A532B), the BMBF-funded Medical Informatics Initiative (HiGHmed, 01ZZ1802A–01ZZ1802Z), the BMBF-funded projects 01IK20337 (RECAST) and 82DZL0098B1, the German Federal Ministry of Research, Technology and Space (82DZL009C1 and 01GL2401A), and the German Research Foundation (Deutsche Forschungsgemeinschaft, DFG)-funded CRC-TR 84 B08, CRC-1449 Z01, project ID 272983813-TRR179, the DFG COVID-19 focus funding project BI1693/2-1, CRC 1449 (project number 431232613), and EXC 3118/1 (project number 533770413).

## AUTHOR CONTRIBUTIONS

I.L., and S.T. conceived, designed, and supervised the project. S.M., A.S., M.Me., J.Lo., R.L.C. and J.Li. performed experiments. S.M. analyzed the data. MA.M., J.R., S.W., B.S., and L.E.S. provided the human specimens and clinical annotation of the patients. MA.M., R.E., and C.C. contributed with discussion of the results. F.T., provided technical and experimental support. S.M., S.T., and I.L., wrote the manuscript; all authors read, revised, and approved the manuscript.

## DATA AVAILABILITY

The generated scRNA-seq data are publicly available via FigShare [DOI/link provided upon publication], and raw sequencing files are deposited at the European Genome-phenome Archive (EGA) under accession number [accession code provided upon publication].

---

**Extended Data Figure 1:**
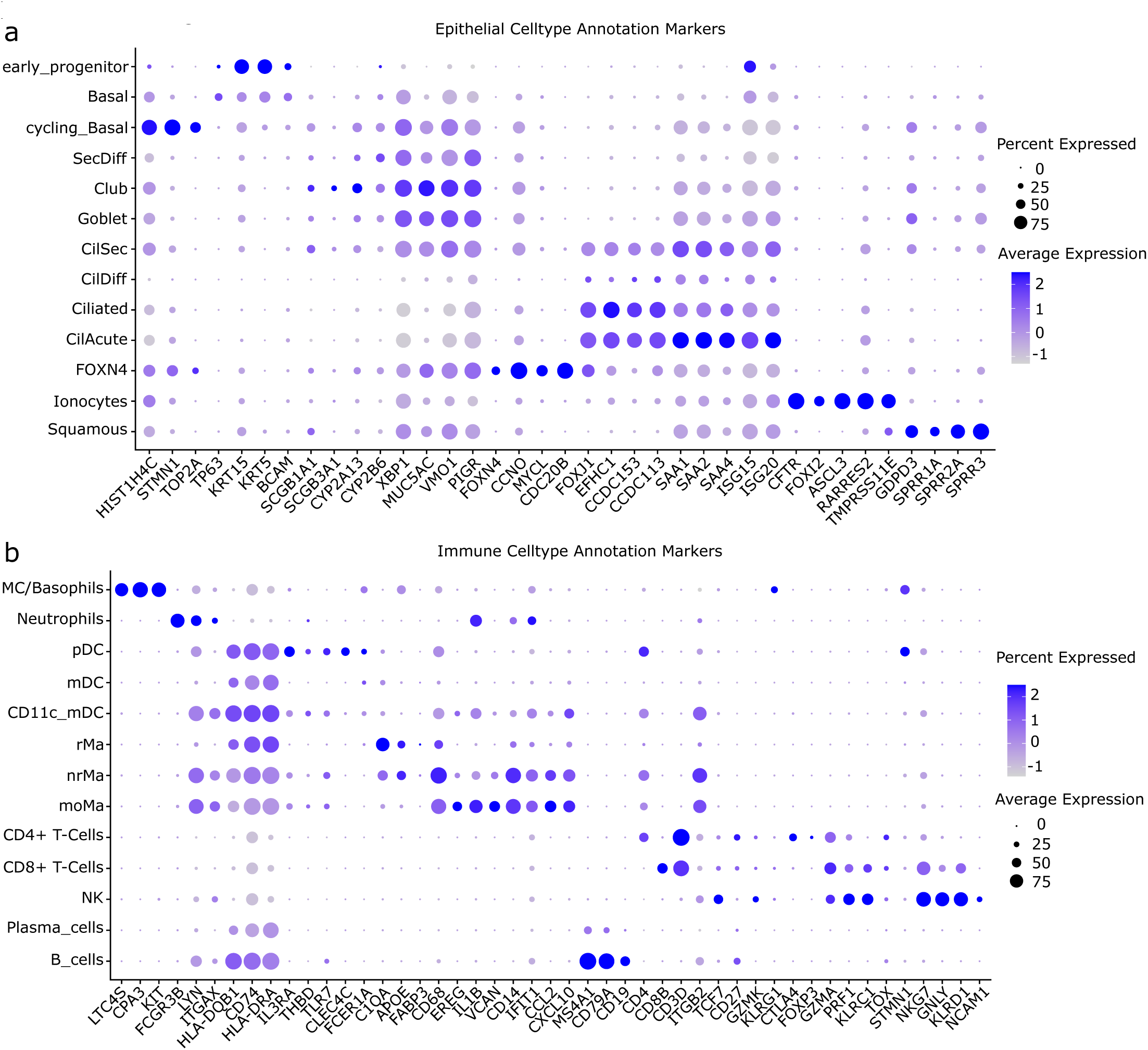
Expression of Identifying Markers Across Cell Types. Mean normalized gene expression of identifying markers shown as percentage of cells expressing each marker and average expression level across all (a) epithelial and (b) immune cells per cell type. All samples were included for cell type annotation (n=68).

**Extended Data Figure 2:**
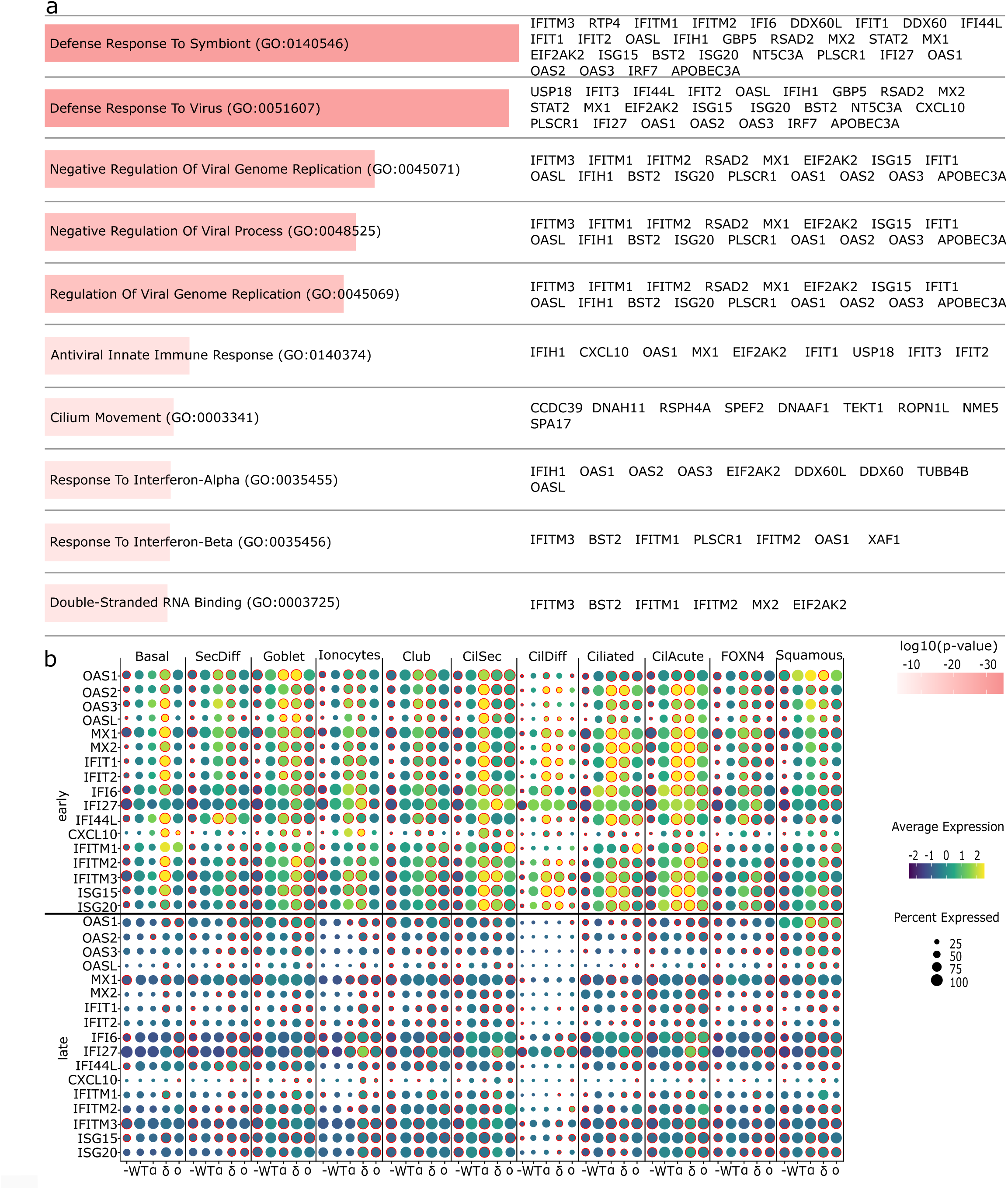
Top 10 Gene Ontology (GO) Terms and corresponding Genes in Epithelium. (a) Top 10 GO Terms resulting from enrichment of top 200 DEGs between controls and COVID-19 patients in epithelium. Overlap of top 200 genes and GO terms are shown on the right. All samples included (n=68). (b) Mean normalized gene expression of interferon stimulated genes shown as percentage of cells expressing each marker and average expression level across epithelial cells. Red circle indicates significance vs. wild-type (fdr < 0.05 & log2FC of >0.1 | < -0.1). Asymptomatic and those without dps information were excluded (n=64).

**Extended Data Figure 3:**
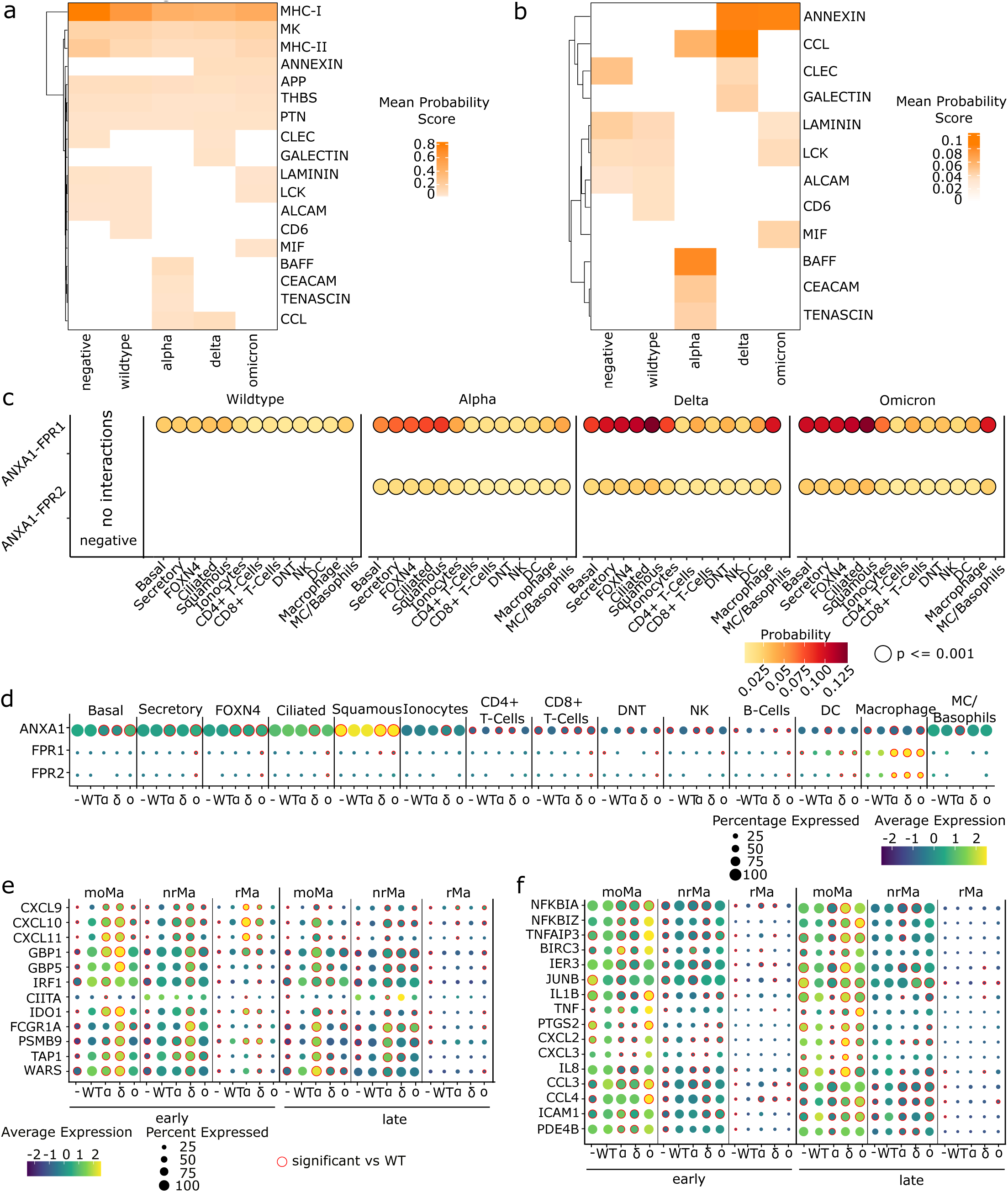
CellChat Pathway Identification and Immune Cell Ligand/Receptor Expression. Mean probability score for (a) the union of the top 10 pathways per condition and (b) pathways with differential probability between conditions (p<0.05).(c) Interaction probability between ligands and receptors in macrophage-targeting ANNEXIN signalling, source celltypes shown on the x-axis (p<0.01). Mean normalized gene expression of (d) ANNEXIN ligands and receptors across variants (e) interferon-related macrophage activation markers and (f) cytokine-response genes. In (d-f), red circles indicate significant differential expression vs. wild-type (fdr < 0.05 & |log2FC| > 1). Panels (a-d) excluded severe samples (n=65). Panels (e-f) excluded severe, asymptomaticpatients and those without dps information (n=61).

**Extended Data Figure 4:**
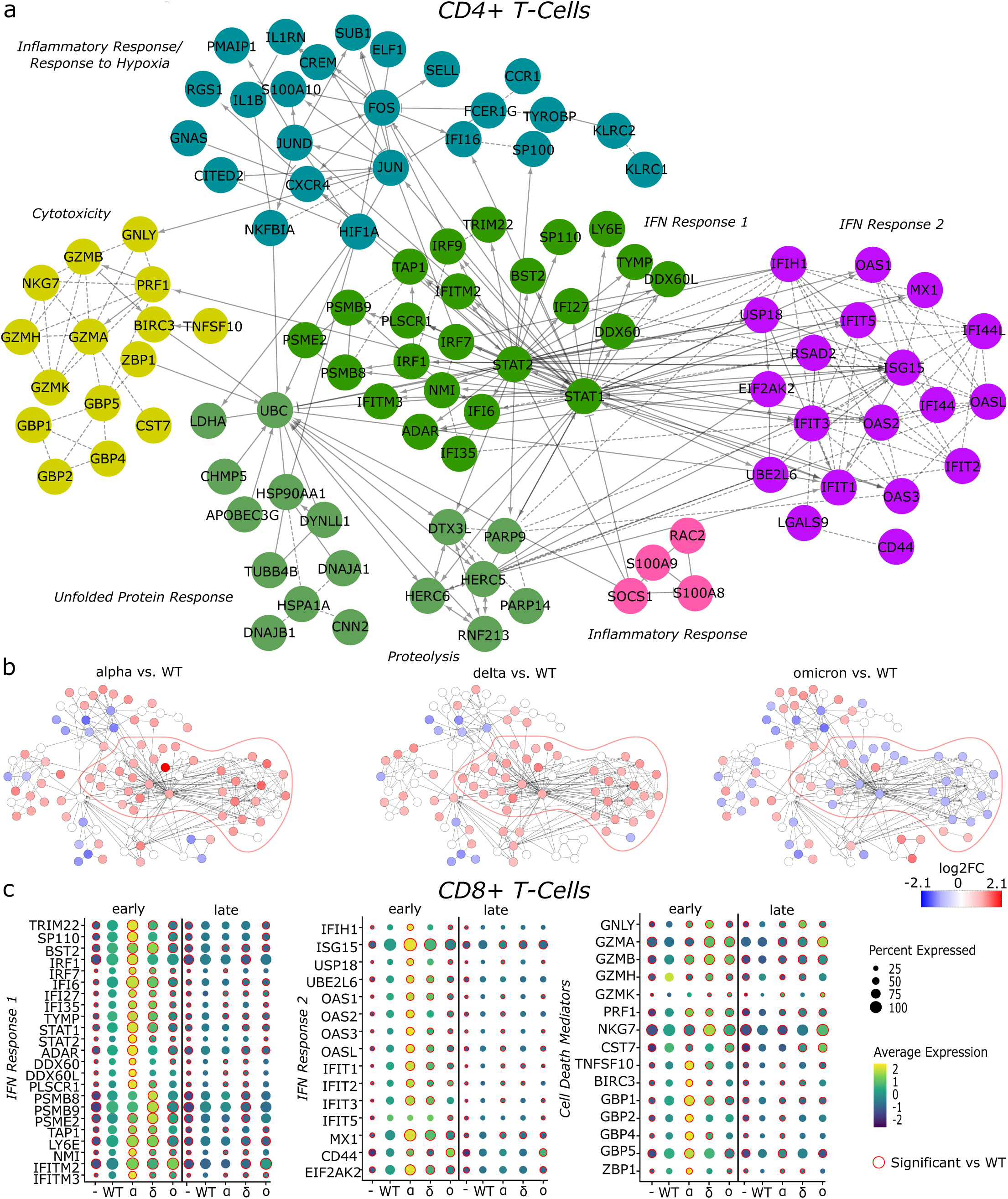
Further Cytoscape Analyses in CD4+ and CD8+ T-Cells. Cytoscape network based on the top 200 DEGs between controls and SARS-CoV-2+ samples in CD4+ T-Cells. Node colour represents (a) module annotation (b) log2FC for each variant vs. WT (fdr < 0.05). A red border marks IFN Response modules. Controls, mild and moderate COVID-19 samples (n=62) were included. (c) Comparison of mean normalized gene expression of genes comprising Interferon Response 1, Interferon Response 2 and the Cell Death Mediator Modules across variants. Red circles indicate significant differential expression vs. wild-type (fdr < 0.05 & |log2FC| > 1). Panel (c) included controls, mild and moderate COVID-19 samples, with dps information (n=61).

## References

1. Loske, J., et al. Pre-activated antiviral innate immunity in the upper airways controls early SARS-CoV-2 infection in children. Nat Biotechnol 40, 319–324 (2022).

2. Wimmers, F., et al. Multi-omics analysis of mucosal and systemic immunity to SARS-CoV-2 after birth. Cell 186, 4632–4651 e4623 (2023).

3. Walsh, J.M.L., et al. Variants and vaccines impact nasal immunity over three waves of SARS-CoV-2. Nat Immunol 26, 294–307 (2025).

4. Mick, E., et al. Upper airway gene expression shows a more robust adaptive immune response to SARS-CoV-2 in children. Nat Commun 13, 3937 (2022).

5. Yoshida, M., et al. Local and systemic responses to SARS-CoV-2 infection in children and adults. Nature 602, 321–327 (2022).

6. Lindeboom, R.G.H., et al. Human SARS-CoV-2 challenge uncovers local and systemic response dynamics. Nature 631, 189–198 (2024).

7. Tilley, A.E., Walters, M.S., Shaykhiev, R. & Crystal, R.G. Cilia dysfunction in lung disease. Annu Rev Physiol 77, 379–406 (2015).

8. Horani, A., Ferkol, T.W., Dutcher, S.K. & Brody, S.L. Genetics and biology of primary ciliary dyskinesia. Paediatr Respir Rev 18, 18–24 (2016).

9. Messingschlager, M., et al. DNA methylation changes during acute COVID-19 are associated with long-term transcriptional dysregulation in patients’ airway epithelial cells. EMBO Mol Med 17, 923–937 (2025).

10. Chang, J.J., et al. Uncovering strain- and age-dependent innate immune responses to SARS-CoV-2 infection in air-liquid-interface cultured nasal epithelia. iScience 27, 110009 (2024).

11. Reuschl, A.K., et al. Evolution of enhanced innate immune suppression by SARS-CoV-2 Omicron subvariants. Nat Microbiol 9, 451–463 (2024).

12. Thorne, L.G., et al. Evolution of enhanced innate immune evasion by SARS-CoV-2. Nature 602, 487–495 (2022).

13. Zhu, Y., et al. Ancestral SARS-CoV-2, but not Omicron, replicates less efficiently in primary pediatric nasal epithelial cells. PLoS Biol 20, e3001728 (2022).

14. Winnett, A.V., et al. Early Mucosal Type II Interferon Limits SARS-CoV-2 Replication in Humans. medRxiv (2026).

15. Jin, S., et al. Inference and analysis of cell-cell communication using CellChat. Nat Commun 12, 1088 (2021).

16. Gemperle, C., et al. Regulation of the formyl peptide receptor 1 (FPR1) gene in primary human macrophages. PLoS One 7, e50195 (2012).

17. Schloer, S., et al. The annexin A1/FPR2 signaling axis expands alveolar macrophages, limits viral replication, and attenuates pathogenesis in the murine influenza A virus infection model. FASEB J 33, 12188–12199 (2019).

18. McArthur, S., et al. Annexin A1 drives macrophage skewing to accelerate muscle regeneration through AMPK activation. J Clin Invest 130, 1156–1167 (2020).

19. Waddell, S.J., et al. Dissecting interferon-induced transcriptional programs in human peripheral blood cells. PLoS One 5, e9753 (2010).

20. Zhang, F., et al. IFN-gamma and TNF-alpha drive a CXCL10+ CCL2+ macrophage phenotype expanded in severe COVID-19 lungs and inflammatory diseases with tissue inflammation. Genome Med 13, 64 (2021).

21. Lama, L., Morozov, P., Garzia, A. & Tuschl, T. Dissection of innate-immune-ligand- and interferon-protein-mediated transcriptional responses in human THP1 cell states. Commun Biol 9, 239 (2026).

22. Martin, E.W., Pacholewska, A., Patel, H., Dashora, H. & Sung, M.H. Integrative analysis suggests cell type-specific decoding of NF-kappaB dynamics. Sci Signal 13(2020).

23. Sharif, O., Bolshakov, V.N., Raines, S., Newham, P. & Perkins, N.D. Transcriptional profiling of the LPS induced NF-kappaB response in macrophages. BMC Immunol 8, 1 (2007).

24. Chua, R.L., et al. COVID-19 severity correlates with airway epithelium-immune cell interactions identified by single-cell analysis. Nat Biotechnol 38, 970–979 (2020).

25. Trump, S., et al. Hypertension delays viral clearance and exacerbates airway hyperinflammation in patients with COVID-19. Nat Biotechnol 39, 705–716 (2021).

26. Zella, D., et al. Interferon-gamma increases expression of chemokine receptors CCR1, CCR3, and CCR5, but not CXCR4 in monocytoid U937 cells. Blood 91, 4444–4450 (1998).

27. Shannon, P., et al. Cytoscape: a software environment for integrated models of biomolecular interaction networks. Genome Res 13, 2498–2504 (2003).

28. World Health Organisation. Historical working definitions and primary actions for SARS-CoV-2 variants., Vol. 2026 (2023).

29. Aylward, B., Liang, W. & WHO-China-Joint-Mission,. Report of the WHO-China Joint Mission on Coronavirus Disease 2019 (COVID-19). Vol. 2026 (World Health Organisation, 2020).

30. Hao, Y.H., et al. Integrated analysis of multimodal single-cell data. Cell 184, 3573– 3587 (2021).

31. Wolf, F.A., Angerer, P. & Theis, F.J. SCANPY: large-scale single-cell gene expression data analysis. Genome Biol 19 (2018).

32. Korsunsky, I., et al. Fast, sensitive and accurate integration of single-cell data with Harmony. Nat Methods 16, 1289–1296 (2019).

33. Polanski, K., et al. BBKNN: fast batch alignment of single cell transcriptomes. Bioinformatics 36, 964–965 (2020).

